# Integrative Multi-Omics Analysis of Barley Genotypes Shows Differential Salt-Induced Osmotic Barriers and Response Phases Among Rootzones

**DOI:** 10.1101/825059

**Authors:** William Wing Ho Ho, Camilla B. Hill, Monika S. Doblin, Megan C. Shelden, Allison van de Meene, Thusitha Rupasinghe, Antony Bacic, Ute Roessner

**Affiliations:** School of BioSciences, The University of Melbourne, Parkville VIC 3010, Australia; School of Veterinary and Life Sciences, Murdoch University, Murdoch, WA 6150, Australia; La Trobe Institute for Agriculture & Food, Department of Animal, Plant and Soil Science, La Trobe University, Bundoora VIC 3086; ARC Centre of Excellence in Plant Energy Biology, School of Agriculture, Food and Wine, University of Adelaide, Glen Osmond SA 5064, Australia; Metabolomics Australia, The University of Melbourne, Parkville VIC 3010, Australia; Melbourne Integrative Genomics, Schools of Mathematics and Statistics and of BioSciences, The University of Melbourne, Parkville VIC 3010, Australia

## Abstract

The mechanisms underlying rootzone-localised responses to salinity stress during early stage of barley development remains fragmentary and elusive. Here, we performed a comprehensive detection of the multi-root-omes (transcriptomes, metabolomes, lipidomes) of a domesticated barley cultivar (Clipper) and a landrace (Sahara) with seedling root growth maintained and restricted in response to salt stress, respectively. Novel generalized linear models were designed to determine differentially expressed genes (DEG) or abundant metabolites (DAM) specific to salt treatments, genotypes, or rootzones (meristematic Z1, elongation Z2, maturation Z3). Based on pathway over-representation of the DEG and DAM, phenylpropanoid biosynthesis is the most statistically over-represented biological pathways among all salinity responses observed. Together with the histological evidence, an intense salt-induced lignin impregnation was found only at the stelic cell wall of Clipper Z2, comparing to a unique elevation of suberin deposition across Sahara Z2. This suggests two differential salt-induced modulations of apoplastic flow between the genotypes. Based on global correlation network construction of the DEG and DAM, callose deposition that potentially adjusted the symplastic flow in roots was almost independent of salinity in rootzones of Clipper, but was markedly decreased in that of Sahara. Through closer examinations of molecular and hormonal clues, we further demonstrate that the salinity response in rootzones of Clipper were mostly at recovery phase, comparing to Sahara with rootzones retained at quiescence. Taken together, we propose that two distinctive salt tolerance mechanisms could exist in Clipper (growth-sustaining) and Sahara (salt-shielding), providing important clues for improving crop plasticity to cope with the deteriorating global salinization of soil.

## INTRODUCTION

Barley (*Hordeum vulgare* L.) is an essential feed, food and brewing crop, and a model system for temperate cereals. As a glycophyte, barley suffers substantial yield loss when grown under saline conditions, with roots acting as the first sensors and responders [Glenn *et al*., 1999, #3173]. Differential responses at the level of either cell types or developmental zones are part of a strategy for the root to respond and acclimate to environmental changes [Dinneny *et al*., 2008, #76723; Sarabia *et al*., 2018, #42076]. Although a large number of studies have investigated salinity responses of plants at the physiological and molecular level [Shelden *et al*., 2013, #95836; Hill *et al*., 2013, #69673], relatively little is known about the early root zone-specific response to salt stress in barley roots. Integrative ‘omics approaches within large-scale experiments, including genomics, transcriptomics, ionomics, proteomics, and metabolomics, can help decipher the interplay of cellular functions at different levels.

In barley, several initial analyses indicate that different developmental zones within the root respond distinctly to salt stress in tolerant and sensitive genotypes. In a previous study, roots of two barley genotypes showed contrasting early growth responses to salt stress during their seedling development: Clipper, a domesticated cultivar, showed sustained seminal root growth, whereas Sahara, an African landrace, showed decreased seminal root growth [Shelden *et al*., 2013, #95836]. An untargeted metabolomics approach identified early changes in root primary metabolites of both genotypes in response to salt stress using seedling roots sectioned into the meristematic (Z1), elongation (Z2), and maturation (Z3) zones [Shelden *et al*., 2016, #60510]. This initial study showed that the processes involved in growth adaptation and coordination of metabolic pathways in barley roots were under spatial and temporal control. In a subsequent study, two *de novo* transcriptome assemblies of Clipper and Sahara were constructed and generalized linear models (GLM) were applied to access spatial, treatment-related, and genotype-specific gene responses along the developmental gradient of barley roots [Hill *et al*., 2016, #53060]. A gradual transition from transcripts related to sugar-mediated signalling at Z1 to those involved in cell wall metabolism in Z2 was observed.

To take advantage of the latest version of the barley reference genome (Morex v2) with increased sequencing depth and genome coverage [Mascher *et al*., 2017, #46618], here we built on our previous work [Hill *et al*., 2016, #53060] and re-visited the twelve transcriptomes using an advanced bioinformatics pipeline with improved gapped-read mapping and functional annotation of genes. To obtain further molecular insights into the impact of salinity at the metabolite level, we adopted a combined targeted metabolomics and lipidomics approach to quantitatively determine the alteration of the corresponding primary metabolites and lipids. Here, we designed a new GLM-based analysis approach to identify the treatment-, genotype-, and root zone-specific differentially expressed genes (DEG) concurrently with the differentially abundant metabolites and lipids (DAM) in barley root zones upon salt stress. Integrated pathway over-representation of the DEG and DAM showed that the salt treatment led to two differential modulations of phenylpropanoid biosynthesis, which likely contributed to the salinity-induced localization changes of cell wall components, such as lignin and suberin, in Clipper and Sahara. As a proof of concept, we further explored the interconnections between affected metabolites and gene expression pathways by construction of global coexpression-correlation networks specific to each barley genotype. Based on our system-wide exploration, we demonstrate that seedlings of both Clipper and Sahara respond to salinity stress differentially, suggesting the distinctive dynamics underpinning the plasticity of different barley genotypes in response to salt stress.

## RESULTS

### Barley Transcriptome-Processing and -Annotation Pipeline

The improved workflow for transcriptome sequence pre-processing, pre-mapping, mapping and transcript analyses is presented in Fig. 1a. We achieved an average mapping efficiency of 95.7 ± 1.6% for the 192 sequenced libraries used in this study (Supplemental Fig. 1a), demonstrating a high degree of sequence conservation among Morex, Clipper, and Sahara at the transcript level. In total, 247,281 out of the 333,926 predicted transcripts (74.1%) of the Morex v2 genome were functionally annotated compared to around 37.4% and 40.1% annotation obtained for the *de novo* assemblies of Clipper and Sahara, respectively [Hill *et al*., 2016, #53060] (Supplemental Fig. 1b). From this we constructed a new counting matrix comprised of the trimmed mean of M-values (TMM)-normalized counts per million (CPM) reads for the twelve transcriptomes (Supplemental Data Set 1).

**Figure 1.**
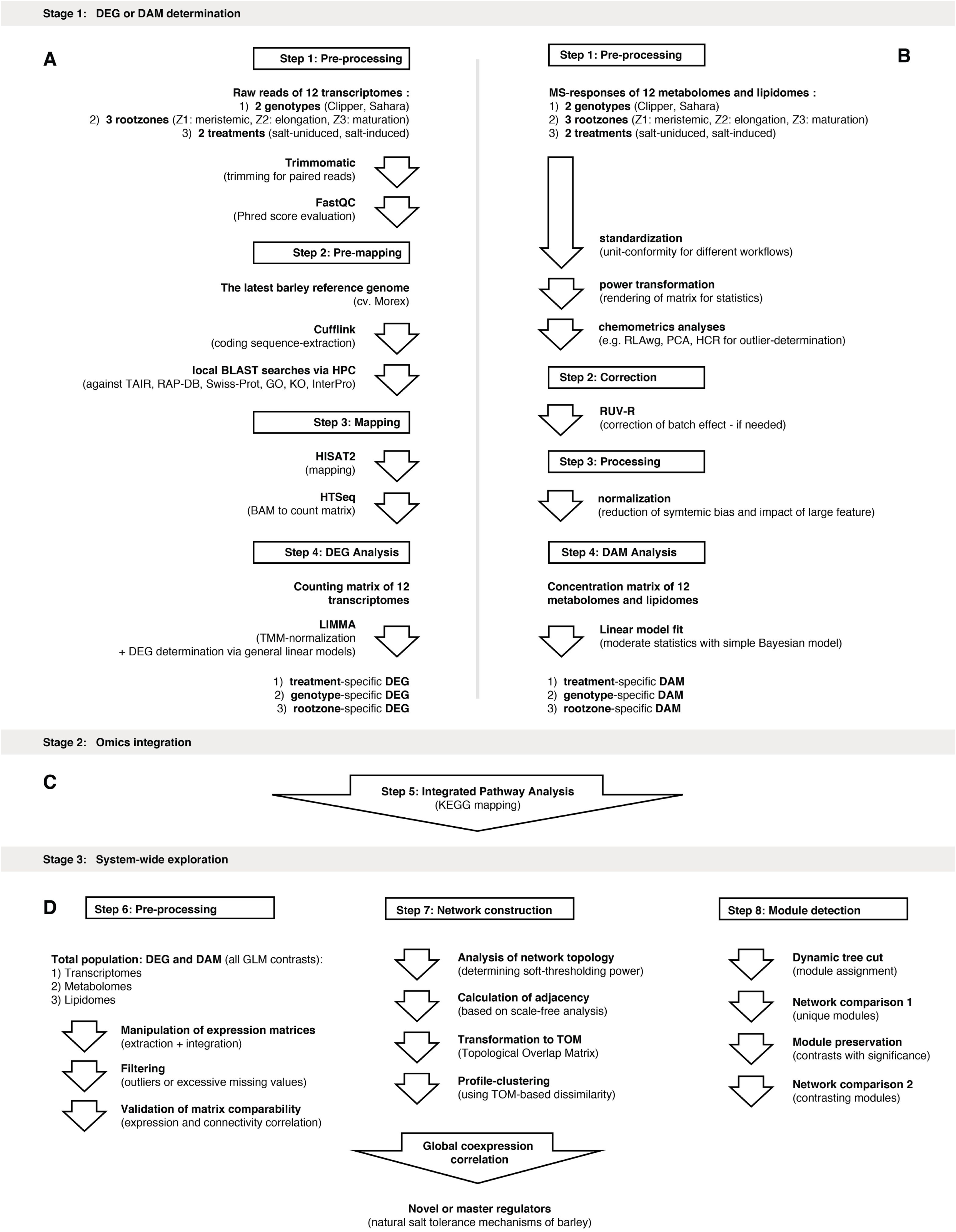
Overview of the bioinformatics pipelines implemented in this study. DAM, differentially abundant metabolite; DEG, differentially expressed genes; GLM, general linear model; GO, Gene Ontology; HCR, hierarchical clustering; HPC, high performance computation; KO, Kyoto Encyclopedia of Genes and Genomes Ontology; MS, Mass Spectrometry; PCA, principal component analysis; RAP-DB, Rice Annotation Project - Database; RLAwg, within-group relative log adjustment; TAIR, The Arabidopsis Information Resource; TMM, trimmed mean normalization; Z1, zone 1 (meristematic zone); Z2, zone 2 (elongation zone); Z3, zone 3 (maturation zone).

### Effects of Salinity on Barley Transcriptomes

We determined the DEG specific to treatment (0mM or 100mM NaCl), genotype (Clipper or Sahara), and root zone (meristematic (Z1), elongation (Z2), or maturation (Z3)). Here, specific GLMs taking the interactions among three factors, namely treatments, genotypes, and root zones into account, were applied to determine genotype- and root zone-specific DEG. Notably, for explaining a phenotype specific to either a particular genotype or root zone, two possibilities exist: differences could either be due to the effect of DEG unique to a genotype or root zone (Fig. 2a), or of DEG common to both genotypes and root zones, but with significant differences in expression (Fig. 2b). To this end, both uniqueness and significance of difference in expression were addressed through specific GLM designs as described in Supplemental Note 1 and the results were summarised in Fig. 2c-f.

**Figure 2.**
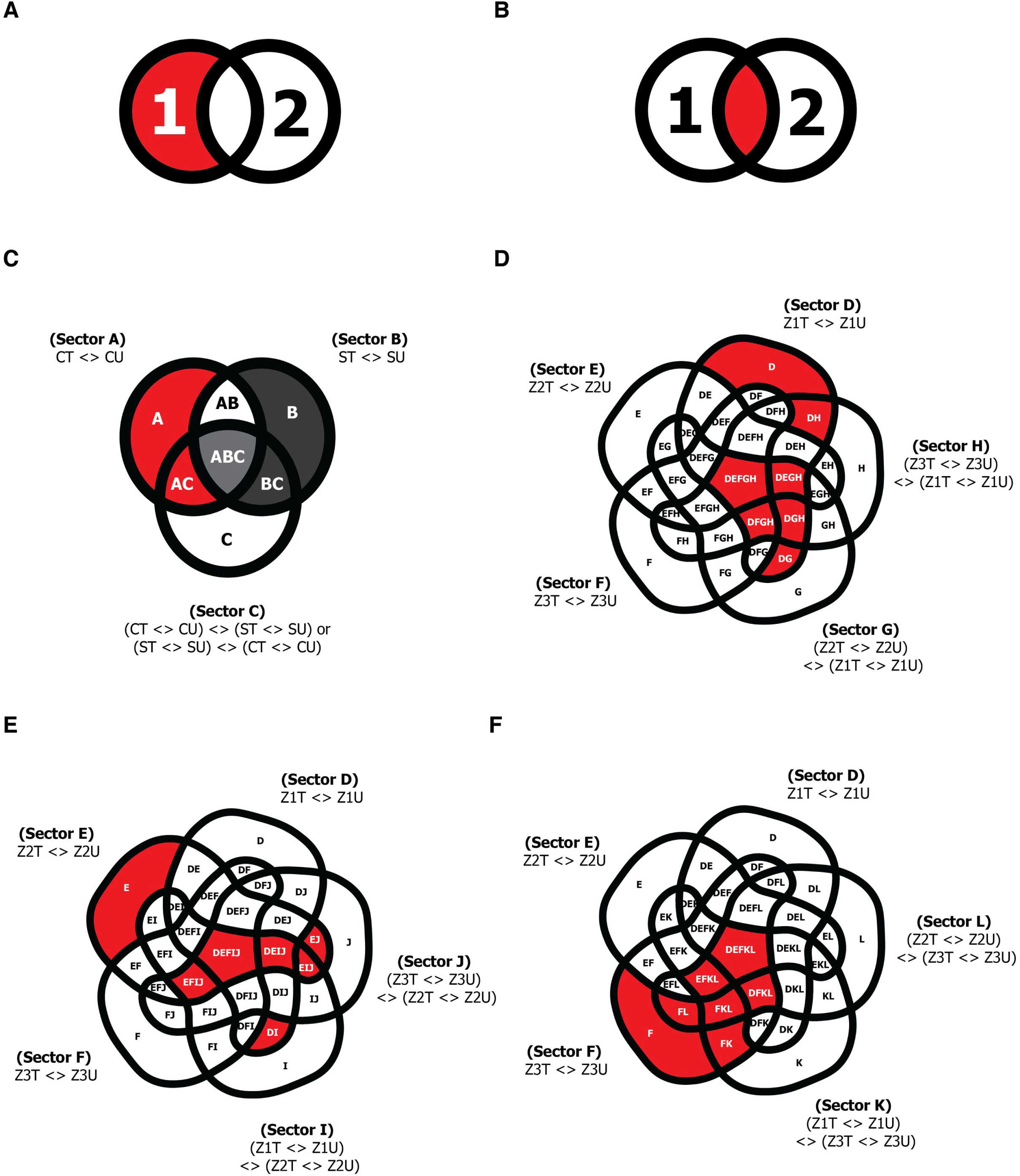
Design of generalized linear models and subsetting for the DEG or DAM determination. Two possible sets of DEG or DAM that could account for the genotype- or root zone-specific phenotypes (colored in red): **(A)** DEG or DAM unique to one genotype or root zone; **(B)** DEG or DAM common to both genotypes or root zones, but showed significant contrast in expression or abundance between the two. Number 1 and 2 in the figures denote two sets of DEG or DAM from two different genotypes/ root zones in comparison. **(C)** Genotype-specific DEG or DAM for each root zone. Subsectors correspond to the Clipper-specific DEG/DAM (including subsectors A, AC) and Sahara-specific DEG or DAM (including subsectors B, BC) in each root zone are highlighted in red and dark grey, respectively. Subsector ABC are common to both Clipper and Sahara (colored in light grey), but defined by GLM contrast in opposite directions: (CT <> CU) <> (ST <> SU), and (ST <> SU) <> (CT <> CU), respectively. **(D-F)** Root zone-specific DEG or DAM of Clipper/ Sahara at **(D)** meristematic zone (Z1), **(E)** elongation zone (Z2), and **(F)** maturation zone (Z3) respectively, and with the corresponding subsectors highlighted in red. CT, salt-treated Clipper; CU, untreated Clipper; DEG, differentially expressed genes; DAM, differentially abundant metabolites; ST, salt-treated Sahara; SU, untreated Sahara; Z1T, salt-treated Z1; Z1U, untreated Z1; Z2T, salt-treated Z2; Z2U, untreated Z2; Z3T, salt-treated Z3; Z3U, untreated Z3; <>, contrast of GLM.

Among the 11,631 detected transcripts (Supplemental Fig. 2a), the GLM-based differential analyses revealed that the abundance of 3,801 transcripts (32.7%) changed significantly (up- or down-regulated) after the salt treatment in Clipper (see Supplemental Data Set 3 for annotated DEG lists). In Sahara, 4,789 transcripts (41.2%) were significantly different in abundance in response to salt treatment relative to their controls, indicating the overall change in gene expression induced by salt was more pronounced in Sahara than in Clipper. From the perspective of root zones within both barley genotypes, 2,148 (18.5%), 4,759 (40.9%), and 3,948 (33.9%) transcripts within the 11,631 quantifiable pool altered significantly after the salt treatment in Z1, Z2, and Z3, respectively. This suggests that the effect of salinity was more substantial in Z2, followed by Z3 and then Z1 at the transcript level in both genotypes. Further, among the 3,801 treatment-specific DEG in |Clipper, 2,774 transcripts (73.0%) were shown to be highly specific to their genotype, compared to 4,144 (86.5%) transcripts within the 4789 treatment-specific DEG in Sahara. In contrast, among the treatment-specific DEG in Z1 (2,148), Z2 (4,759), and Z3 (3,948), only 1,225 (57.0%), 2,710 (56.9%), and 1,501 (38.0%) transcripts were found to be highly specific to their respective root zones. This indicates the salt-induced responses at transcript level in roots were more dependent on their genotypes than on their developmental zones.

To determine which biological processes are most prominent in the two genotypes upon salt-stress, treatment-specific DEG in each genotype were classified into seven groups according to their spatial distribution in barley roots. Each group was then subject to enrichment analysis of gene ontology (GO) with a focus on the category of biological processes (Fig. 3; Supplemental Data Set 5).

**Figure 3.**
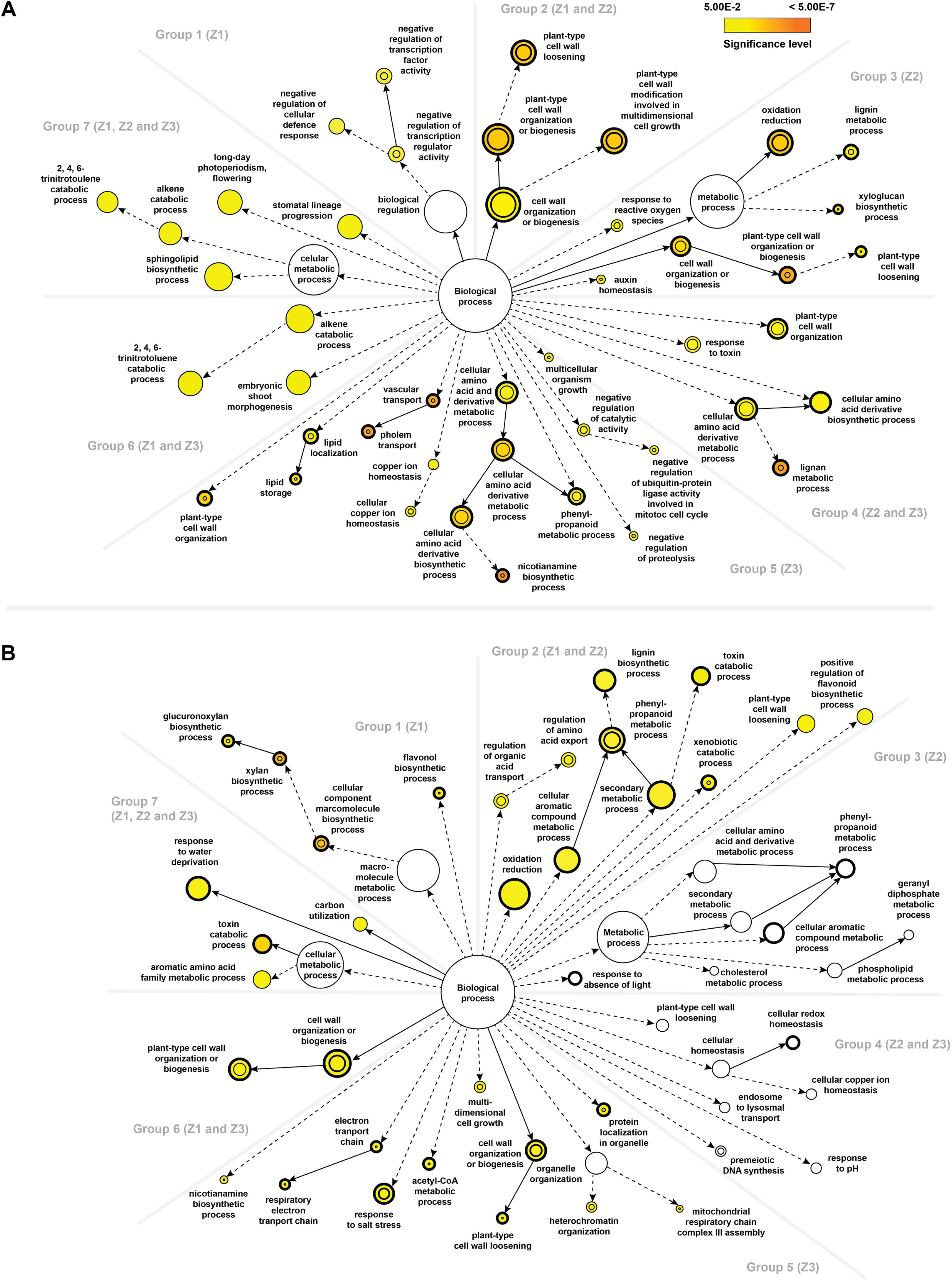
Comparisons of the statistical over-representation of GO categories between different root zones of the two barley genotypes upon salt stress. **(A)** Statistically over-represented GO categories unique to or shared between different root zones of Clipper. **(B)** Statistically over-represented GO categories unique to or shared between different root zones of Sahara. Nodes represent GO categories and node-size is proportional to the number of detected genes for each node. Categories under the same GO hierarchy are linked by interconnected arrows (known as edges) and intensity of node-color indicates the significance level of statistical overrepresentation determined by Fisher’s exact test with adjusted p value <0.05 as cutoff as per legend. For reference only, a threshold of 0.2 is set for those sectors showing no significant over-representation and white-colored nodes are used to visualize those ontologies closed to the threshold. Dotted edges indicate one or more hierarchies of GO, which have no statistical significance in the over-representation test and are determined as redundant via REVIGO, were not shown for clarity. DEG statistically over-represented in both treatment-specific and genotype-specific analyses are denoted by node with thickened outlines. DEG statistically over-represented in both treatment-specific and root-zone-specific batches are denoted by inner circle of nodes.

**Group 1** comprised of DEG found only in Z1. GO analysis via BiNGO [Maere *et al*., 2005, #47202] and REVIGO [Supek *et al*., 2011, #81163] revealed the most significant over- representation of this group were regulation of transcription and cellular defense response genes in Clipper, and biosynthesis of hemicelluloses including xylan and its derivatives in Sahara. **Group 2** included DEG found in both Z1 and Z2. GO analysis indicated the genes to be mostly enriched in cell wall modification (in particular cell wall loosening) for Clipper, and phenylpropanoid metabolism for Sahara. **Group 3** consisted of DEG found only in Z2. While plant-type cell wall organization as well as lignin metabolism genes were strongly over-represented in Clipper, no significant enrichment of any GO category could be detected in Sahara. **Group 4** represented DEG found in Z2 and Z3. Lignan metabolism genes and related processes were drastically enriched in Clipper, but similar to Group 3, no significant over-representation was detected in Sahara. **Group 5** consist of DEG found only in Z3. In Clipper, nicotianamine metabolic process as well as vascular transport genes were ranked top in the overrepresentation list, compared to the enrichment of genes encoding proteins targeted to the mitochondrion, response to salt stress, and cell wall organization (xyloglucan metabolism) in Sahara. **Group 6** represented DEG found in both Z1 and Z3. This cluster was enriched in trinitrotoluene catabolism and related processes for Clipper, cell wall organization or biosynthesis for Sahara. **Group 7** contains DEG found in all three root zones. Sphingolipid biosynthesis genes were enriched in Clipper, whereas toxin metabolism was the most significantly overrepresented GO category in Sahara.

### Effects of Salinity on the Barley Metabolomes and Lipidomes

Next, we performed quantitative metabolomics and lipidomics analyses in the same root tissue samples in order to provide a complementary perspective to the early salt responses of barley seedling roots. A total of 154 compounds (22 sugars or sugar alcohols, 15 small organic acids, 32 amines or amino acids, 18 fatty acids, and 67 lipids) were quantified using four mass spectrometry-based metabolomics and lipidomics methods (Supplemental Data Set 2). The bioinformatics pipeline for elucidating the treatment, genotype, and root zone-specific DAM is illustrated in Fig. 1b (see Supplemental Data Set 4 for annotated DAM lists). Notably, GLM used in the DAM determination here were identical to that of the transcriptomic analyses to facilitate the subsequent omics comparisons and integration. The GLM-based differential analyses showed that the abundance of 82 (53.3%) and 61 compounds (39.6%) varied significantly with salt treatment in both Clipper and Sahara relative to their controls, respectively (Supplemental Fig. 2b). Across the root-zones, the abundance of 66 (42.9%), 31 (20.1%), and 30 (19.5%) compounds among the 154 quantifiable pools of metabolites changed significantly after the salt treatment in Z1, Z2, and Z3 of both genotypes, respectively.

Furthermore, within the 82 treatment-specific DAM in Clipper, 55 compounds (67.1%) were shown to be highly specific to this genotype, compared to 42 compounds (68.9%) among the 61 treatment-specific DAM in Sahara. In comparison, among the treatment-specific DAM in Z1 (66 compounds), Z2 (31 compounds), and Z3 (30 compounds), 62 (93.9%), 26 (83.8%), and 21 (70.0%) compounds were found to be highly specific to their respective root zones. The differential analyses at the primary metabolite and lipid levels suggest a higher degree of dependence of the salt-induced responses on root-zones than on genotype, compared to the transcriptional level. This implies an intriguing dynamic, where gene expression differences due to salt treatment are dominated by genotype and the downstream metabolic outcome is more influenced by root zones.

To provide insight as to which metabolic groups are most markedly different between the two genotypes upon salt stress, treatment-specific DAM in each genotype were classified into seven groups according to their spatial distribution in barley roots (Fig. 4). Each group was then subject to metabolite set enrichment analysis (MSEA) via MBROLE2 [López-Ibáñez *et al*., 2016, #73627], with results available in Supplemental Data Set 6.

**Figure 4.**
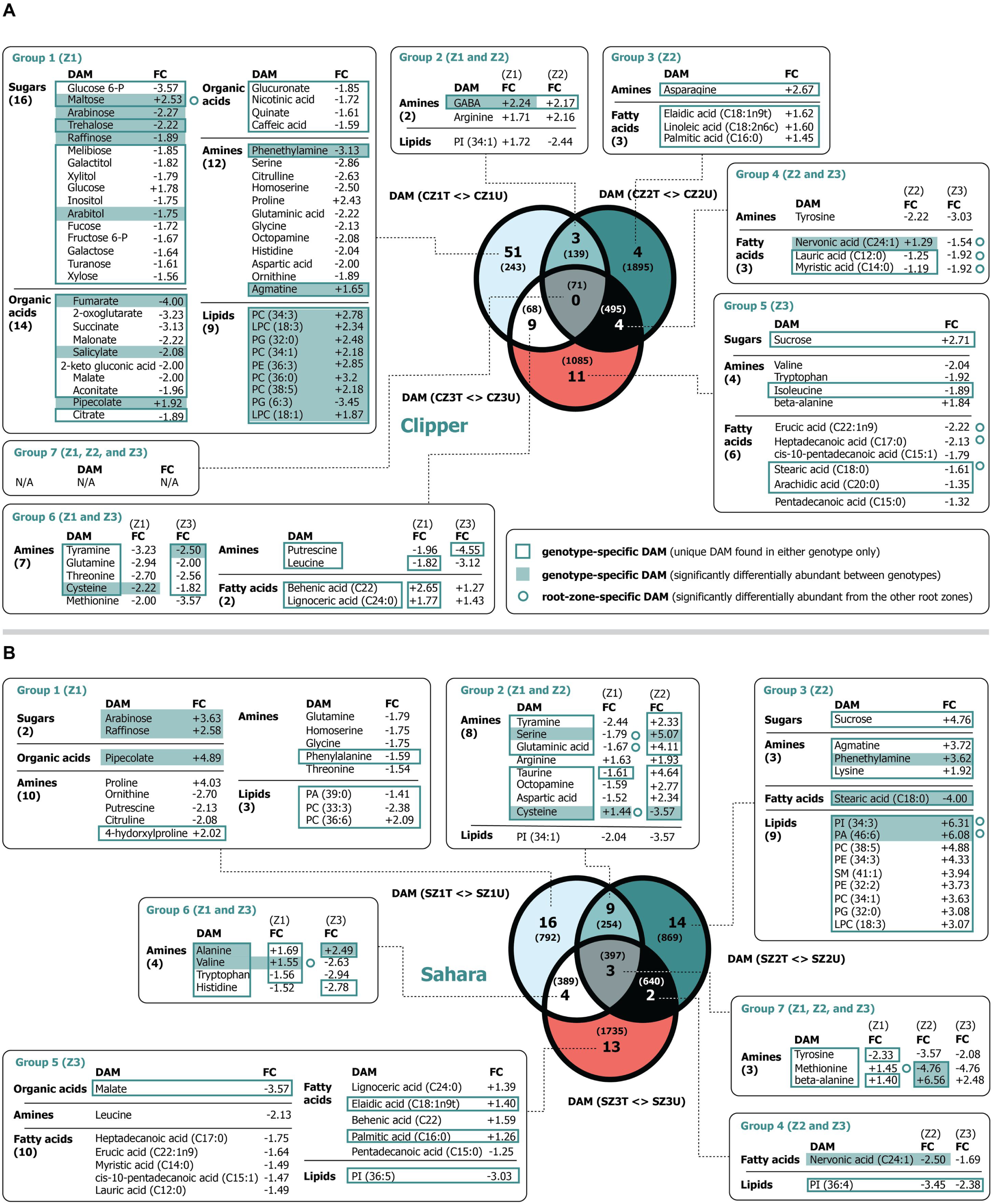
An integrated view of the DAM of the two barley genotypes in response to salt stress. **(A)** DAM unique to or shared among different root zones of Clipper. **(B)** DAM unique to or shared among different root zones of Sahara. Parentheses in each textbox refer to the number of metabolites within each metabolic subgroup. Parentheses in Venn diagrams: corresponding number of DEG in each sector. CZ1T, salt-treated Z1 in Clipper; CZ1U, untreated Z1 in Clipper; CZ2T, salt-treated Z2 in Clipper; CZ2U, untreated Z2in Clipper; CZ3T, salt-treated Z3 in Clipper; CZ3U, untreated Z3in Clipper; DAM, differentially abundant metabolites; DEG, differentially expressed genes; FC, fold change; LPC, lysophosphatidylcholine; PA, phosphtatidic acids; PC, phosphtati-dylcholines; PE, phosphtatidylethanolamines; PG, phosphtatidylglycerols; PI, phosphtatidylinositols; PS, phosphtatidylserines; SM, sphingomyelins; SZ1T, salt-treated Z1 in Sahara; SZ1U, untreated Z1 in Sahara; SZ2T, salt-treated Z2 in Sahara; SZ2U, untreated Z2 in Sahara; SZ3T, salt-treated Z3 in Sahara; SZ3U, untreated Z3 in Sahara; Z1, zone 1 (meristematic zone), Z2, zone 2 (elongation zone); Z3, zone 3 (maturation zone); -P, phosphate.

In brief, **Group 1** comprised of DAM found only in Z1. MSEA revealed the most significant enrichment was the biosynthesis of alkaloids derived from ornithine, lysine and nicotinic acid in Clipper, and arginine and proline metabolism in Sahara. **Group 2** included DAM found in both Z1 and Z2. While there is no significantly enriched functional role detected in Clipper, metabolites were mostly enriched in aminoacyl-tRNA biosynthesis for Sahara. **Group 3** consisted of DAM in Z2. While the biosynthesis of unsaturated fatty acids was strongly over-represented in Clipper, glycerophospholipid metabolism was enriched in Sahara. Notably, nine lipid candidates were up-regulated up to 6.3 fold in Sahara (Fig. 4b), but not in Clipper. **Group 4** represented DAM found in both Z2 and Z3. Whereas fatty acid biosynthesis was drastically enriched in Clipper, no significant over-representation was detected in Sahara. **Group 5** consist of DAM in Z3. In Clipper, aminoacyl-tRNA biosynthesis, glucosinolate biosynthesis, as well as biosynthesis of unsaturated fatty acids were ranked top in the enrichment list, compared to the biosynthesis of unsaturated fatty acids only in Sahara. **Group 6** represented DAM in both Z1 and Z3. These sectors were mostly enriched in biosynthesis of aminoacyl-tRNA for both Clipper and Sahara. No significantly enriched functional role set could be detected for **Group 7**.

### The Over-Represented Salinity Effect on the Barley Root-Omes

We utilized KEGG mapper to perform an integrated pathway analysis for the three omics datasets (Fig. 1c) [Kanehisa *et al*., 2012, #87880]. According to the number of matched DEG and DAM hits, biological pathways statistically over-represented at transcript and/or primary metabolite level in response to salinity were ranked in descending order, where the biosynthesis of phenylpropanoids (such as monolignols, flavonoids, lignins, and suberins) were enriched at both levels and identified at the top of the list (Supplemental Data Set 7). In order to visualize the specific post-salinity effect on the biosynthesis of phenylpropanoids, we calculated the Z-scores of the TMM-normalized CPM for the transcripts involved and of the normalized concentration for the primary metabolites detected. In addition, we adopted an established method to perform a detailed quantification of the phenylpropanoid contents across our root samples (Supplemental Data Set 2) [Vanholme *et al*., 2012, #42742] and computed their Z-scores. The relative abundance of transcripts, primary metabolites, and phenylpropanoids at different root zones of the two barley genotypes were integrated and illustrated in the pathway frameworks modified based on the corresponding KEGG repository (Fig. 5; Supplemental Figs. 3 and 4).

**Figure 5.**
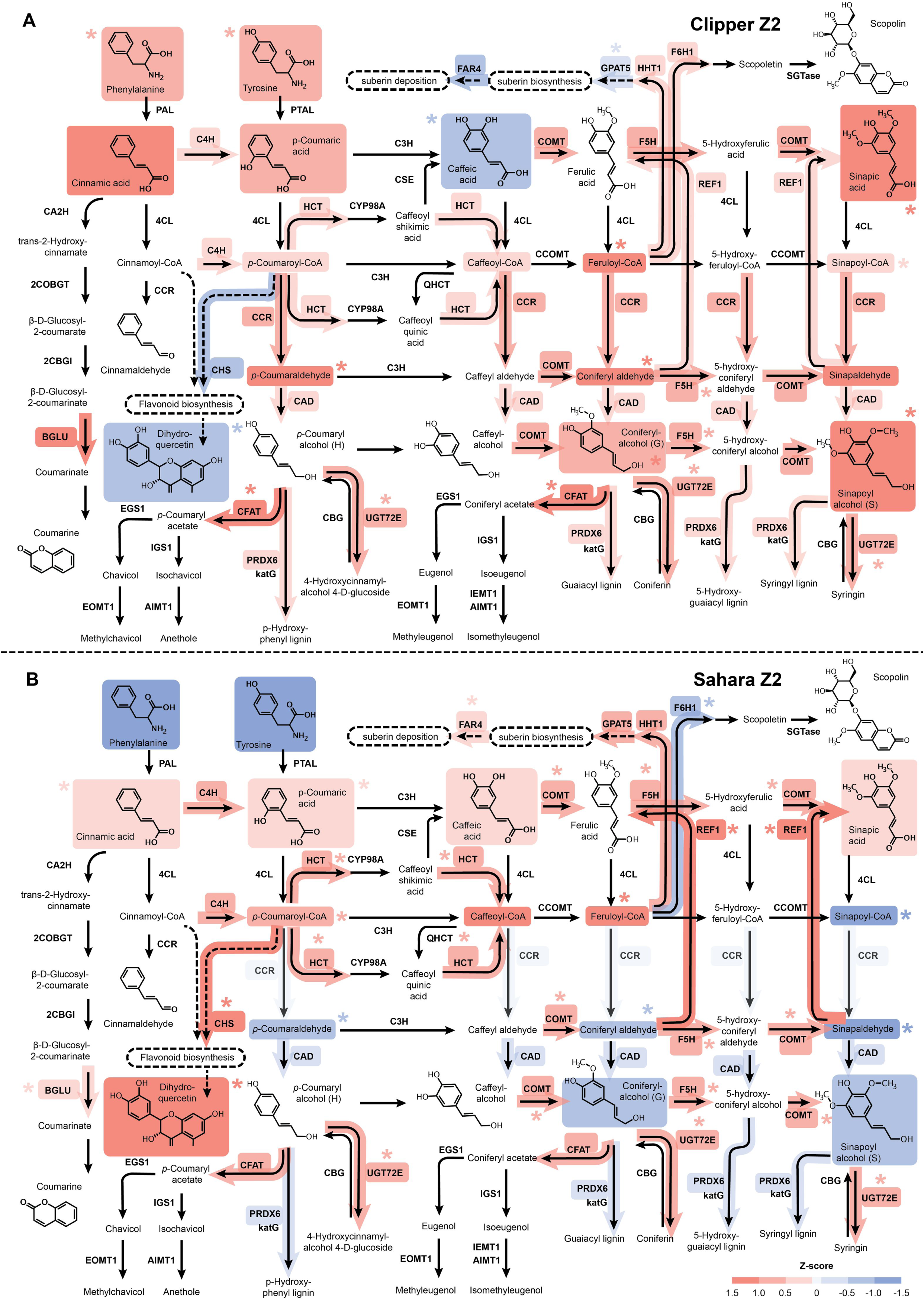
Standardized abundance of transcripts and metabolites involved in phenylpropanoid biosynthesis at the elongation zone (Z2) of the two barley genotypes under salt stress. **(A)** The abundance of transcripts and metabolites involved in the biosynthesis at Clipper Z2. **(B)** The abundance of transcripts and metabolites involved in the biosynthesis at Sahara Z2. Standardized abundances of transcripts and metabolites shown are the Z-scores for TMM-normalized CPM and median-normalized concentration respectively. Level of the standardized abundance (i.e. positive, negative and zero Z-score) is indicated by intensity of shading in red, blue and pale grey, respectively. Asterisks denote statistically significant differentiation of transcript- and metabolite-abundance (with Benjamini-Hochberg adjusted p value < 0.05) after the salt stress compared to the untreated control. Standardized abundance of only the transcripts with significant degree of sequence similarities to the characterized homologs (E-value < 1.00E-3) and the metabolites within the limit of detection of methodologies and instrumentations adopted in this study are shown. Abundance details of these pathway components at different root zones of the two barley genotypes before and after the salt treatment can be found in Supplementary Figure 6 and 7. AIMT1, trans-anol O-methyltransferase; BGLU, beta-glucosidase; CA2H, cinnamic acid 2-hydroxylase; CBG, coniferin beta-glucosidase; 2CBGI, 2-coumarate β-D-glucoside isomerase; CAD, cinnamyl-alcohol dehydrogenase; CCOMT, caffeoyl-CoA O-methyltransferase; CCR, cinnamoyl-CoA reductase; CFAT, coniferyl alcohol acyltransferase; C3H, p-coumarate 3-hydroxylase; C4H, cinnamate 4-hydroxylase; CHS, chalcone synthase; 4CL, 4-coumarate-CoA ligase; COBGT, 2-coumarate O-beta-glucosyltransferase; COMT, caffeic acid 3-O-methyltransferase; CSE, caffeoylshikimate esterase; CYP98A, coumaroylquinate(coumaroylshikimate) 3’-monooxygenase; EGS1, eugenol synthase; EOMT1, eugenol/chavicol O-methyltransferase; F5H, feruiate-5-hydroxylase; F6H1, feruloyl-CoA ortho-hydroxylase; FAR4, fatty acid reductase 4; GPAT5, glycerol-3-phosphate acyltransferase 5; HCT, shikimate O-hydroxycinnamoyltransferase; HHT1, hydroxyacid O-hydroxycinnamoyltransferase 1; IEMT1, (iso)eugenol O-methyltransferase; IGS1, isoeugenol synthase; katG, catalase-peroxidase; PAL, phenylalanine ammonia-lyase; PTAL, phenylalanine/tyrosine ammonia-lyase; PRDX6, peroxiredoxin 6; REF1, coniferyl-aldhyde dehydrogenase; SGTase, scopoletin glucosyltransferase; QHCT, quinate O-hydroxycinnamoyltransferase; UGT72E, coniferyl-alcohol glucosyltransferase

### Salinity-Induced Abundance and Localization Shifts of Phenylpropanoids in the Barley Root Zones

The biosynthetic pathway of phenylpropanoids can be generally divided into three main stages: the general phenylpropanoid pathway from phenylalanine to CoA-esters, monoligonol-specific pathway from CoA-esters to monolignols, and lignin-specific pathway from monolignols to oligolignols or lignin polymers. For Z1 of both barley genotypes (Supplemental Fig. 3), genes involved in all three stages of the biosynthesis remained weakly expressed as in the untreated controls. In line with the detection at the RNA level, negative standardised log_2_ concentration (Z-scores) were recorded for almost all of the metabolic intermediates (Supplemental Figs. 5 and 6). Histochemical staining also showed no observable difference in abundance and localization of phenylpropanoids such as lignin and suberin after salt treatment (Supplemental Figs. 7 and 8), implying that phenylpropanoid production in Z1 was not induced by salt.

**Figure 6.**
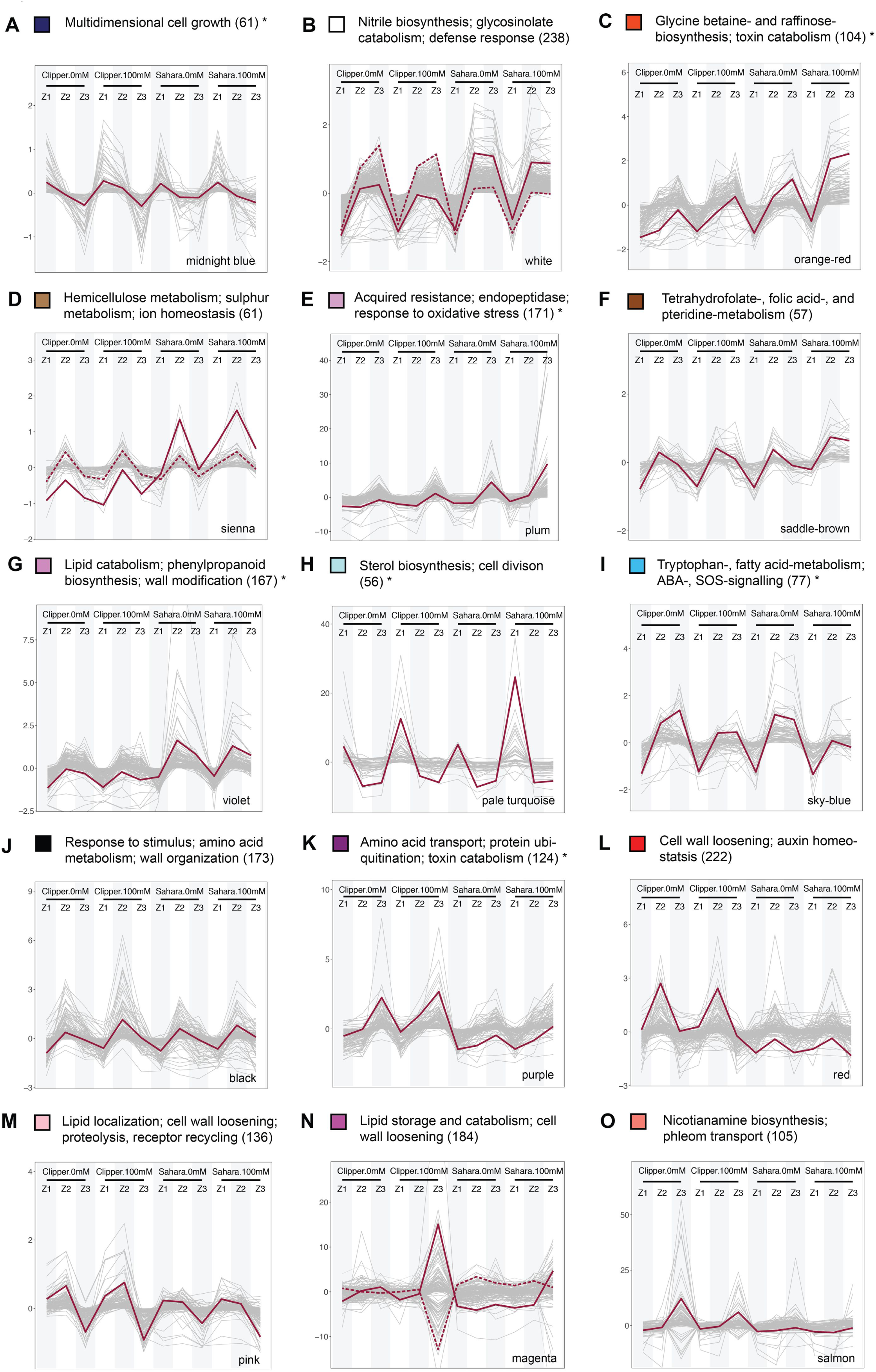
Selected modules of weighted coexpression correlation networks showing abundance profiles of transcripts and metabolites. **(A)** The abundance profile unique to Clipper. **(B-l)** The abundance profiles unique to Sahara. **(J-O)** The abundance profiles significantly contrast between the two barley genotypes. Profiles showing either positive or negative correlations by clustering abundance into differently colored modules through weighted correlation networks. Additional profiles with less obvious differentiation between the two genotypes can be found in **Supplemental** Figure 11. Color of each module consistent with Figure 6. The most representative trend or centroid of each module represented by solid lines are determined by k-mean clustering (distance method: Pearson) with optimal number of clusters calculated from within-group sum of square method **(Madsen and Browning, 2009).** Second the most representative centroid (if any) is indicated by a dotted line. Only expression profiles within 99^th^ percentile are shown for clarity. Annotation of each co-expression clusters are determined by means of the statistical enrichment of GO categories below the cutoff (adjusted *p* value < 0.05) and specific biological role of each module specified here is designated by manual curation of the enrichment outcomes. Asterisks denote the clusters with no significant overrepresentation and anotations assigned to these clusters are the GO categories with the highest possible level of significance **(Supplemental Data Set 8).** Annotated lists of members for each module with significant match (E-value < 1.00E-4) against TAIR10 genome release (version: Jun 2016) ranked in descending order according to kME of members can be found in **Supplemental Data Set 9.**

**Figure 7.**
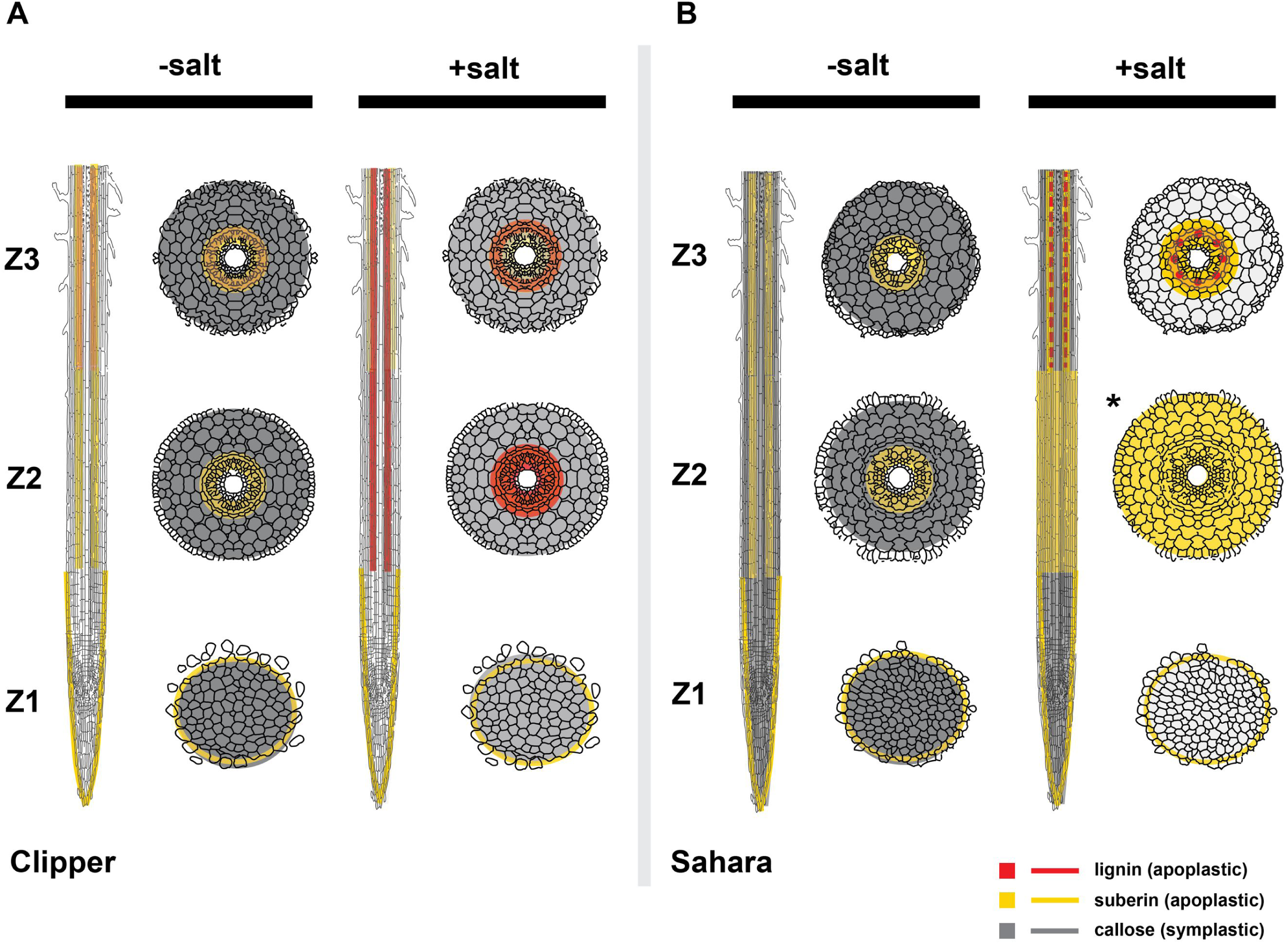
A model of cell wall modifications in roots of Clipper and Sahara in presence or absence of salinity stress. Longitudinal and the corresponding transverse sections of different root zones for **(A)** Clipper and **(B)** Sahara with or without salinity stress. Color Intensities represent the relative levels of lignin, suberin, or callose, which were quantified based on their precursors detection through LC/GC-MS and/or RNA-seq with support of the direct detection of the compounds through histological methods. Localization of each compound was determined by the histo-/immuno-chemical stainings with proven specificity and optical filters applied for minimizing any autoflu-oresence or background signals. Red broken line (Sahara Z3, +salt) represents lignin deposition at only vasculature. Asterisks indicate root zones at cortical, endodermal and/or stelic region with concurrent localizations of suberin and callose (where only suberin is shown for clarity).

In Clipper Z2 (Fig. 5a), transcripts encoded for enzymes involved in the general phenylpropanoid pathway, including: CINNAMATE 4-HYDROXYLASE (C4H), SHIKIMATE O-HYDROXYCINNAMOYLTRANSFERASE (HCT), FERULOYL-COA ORTHO-HYDROXYLASE (F6H1); the monolignol-specific pathway, including: CINNAMOYL-COA REDUCTASE (CCR), CAFFEIC ACID 3-O-METHYLTRANSFERASE (COMT), FERULATE-5-HYDROXYLASE (F5H), CINNAMYL-ALCOHOL DEHYDROGENASE (CAD); and the lignin-specific pathway, including: CONIFERYL ALCOHOL ACYLTRANSFERASE (CFAT), PEROXIREDOXIN6 (PRDX6), CONIFERYL-ALCOHOL GLUCOSYLTRANSFERASE (UGT72E) were either elevated in expression or maintained with positive Z-score after the salt treatment. Consistently, the amount of the detected monolignols, including coniferyl alcohols (guaiacyl (G)-units of lignin) and sinapoyl alcohols (syringyl (S)-units of lignin), were also shown to be significantly induced by salt (Supplemental Fig. 6n,o). The active production of lignins at this root zone was further supported by Basic Fuchsin staining, which indicated a significant increase in lignin impregnation to cellulosic cell walls localized at the outer stelic regions (including endodermis, pericycle, and xylem) of Clipper roots after salt treatment (Supplemental Fig. 7e,f and 7m,n). Further, gene products of *CHALCONE SYNTHASE* (*CHS*) are known to divert intermediates of the general phenylpropanoid pathway for flavonoid production [Heller and Hahlbrock, 1980, #69625]. Weak expression of *CHS* and low levels of flavonoids, such as dihydroquercetin, were consistently detected in Clipper Z2. Notably, transcription of *GLYCEROL-3-PHOSPHATE ACYLTRANSFERASE 5* (*GPAT5*) and *FATTY ACID REDUCTASE 4* (*FAR4*), involved in biosynthesis and deposition of root suberin [Beisson *et al*., 2007, #23927; Domergue *et al*., 2010, #7520], were suppressed by the stress and maintained with negative Z-score, respectively (Supplemental Fig. 5i,j). These data are consistent with the observations of the salinity-induced decline in suberin levels visualized by Fluorol Yellow stain throughout Z2 (Supplemental Fig. 8b,h).

For Sahara Z2 (Fig. 5b), a significant increase of *CHS* expression diverted most phenylpropanoids towards the accumulation of dihydroquercetin (Supplemental Fig. 6p). Together with the low expressions of *CCR* and *CAD*, the accumulation of monolignols and their precursors was restricted and no observable increase of lignin levels in the endodermal region could be detected histochemically after salt stress (Supplemental Fig. 7g,h and o,p). Furthermore, in contrast to Clipper Z2, there was higher abundance of *HYDROXYACID O-HYDROXYCINNAMOYLTRANSFERASE 1* (*HHT1*), *GPAT5* and *FAR4* transcripts in this root zone (Supplemental Fig. 5h-j). The active biosynthesis of suberin inferred from the levels of biosynthetic enzyme transcripts was histologically confirmed with elevated levels of suberin observed in the epidermis and across the subepidermal region of the root zone (Supplemental Fig. 8n,t).

In Z3 of Clipper (Supplemental Fig. 4a), as in Z2, restriction to the production of flavonoids persisted. Salinity-induced accumulation of lignins was also maintained, but found to be limited to G-units and localized at the endodermal and vascular regions (Supplemental Fig. 7a,b). But in contrast to Clipper Z2, an increased biosynthesis and deposition of suberin at the endodermal and stele regions was supported by the consistently higher abundance of *GPAT5*, and *FAR4* transcripts (Supplemental Fig. 5i-j) and by histochemical staining (Supplemental Fig. 8a,g), respectively.

For Z3 of Sahara (Supplemental Fig. 4b), relatively higher transcript abundance of *CCR* and *CAD* compared to Z2 upon salt treatment was detected. Intriguingly, the resulting metabolic changes led to higher accumulation of G-units of lignin and intense deposition of lignin mostly in the xylem vessels of this root zone (Supplemental Fig. 7c,d). Also, in line with the increased Z-score for *CHS* and the suberin-related transcripts (such as *HHT1*, *GPAT5*, and *FAR4*) compared to Clipper Z3 (Supplemental Fig. 5b,h-j), a significant increase of dihydroquercetin and of endodermal and stele suberin deposition was recorded at Sahara Z3, respectively (Supplemental Figs. 6p and 8m,s).

Taken together, the omics datasets at the transcriptional and metabolic levels combined with the histological observations indicate a strong differentiation in biosynthesis and localization of the phenylpropanoids between the two barley genotypes upon salt stress.

### Global Intercorrelations of Salt Stress on the Barley Root-Omes

We next extracted the abundance matrices of transcripts and metabolites that were significantly different in at least one of the GLM-based DEG or DAM determinations in Clipper (3,802 transcripts, 83 metabolites) and Sahara (6,477 transcripts, 61 metabolites). Global co-expression-correlation networks specific to the two barley genotypes were constructed via WGCNA [Langfelder and Horvath, 2008, #26838] (detailed in Methods) to illustrate the system-wide consequences induced by salinity stress (Fig. 1d). In these networks, each “leaf” or short vertical line represents an abundance profile of one transcript, metabolite or lipid. Any interconnected lines within the same “branch” indicate profiles with highly correlated pattern of abundance. Based on the “guilt-by-association” principle as defined in [Saito *et al*., 2008, #659], co-regulated genes and metabolites among each co-expression cluster or “branch” are likely to have common functional roles. To systematically define the “branch”, we applied dynamic tree cut [Langfelder and Horvath, 2008, #26838] to each network and the module assignment was performed to colour code each highly correlated cluster (aka module) (Fig. 6).

For those modules that were unique to either Clipper or Sahara, or were common to both genotypes but significantly contrasted in abundance patterns (Supplemental Table 1), we generated parallel profiles to visualize their variations of abundance in response to salt stress (Fig. 7 and Supplemental Fig. 10). Annotation of each co-expression cluster was determined by means of the statistical enrichment of GO categories below the cutoff (adjusted *p* value ≤ 0.05) and their specific biological roles were designated through manual curation of the enrichment outcomes (Supplemental Data Set 8). Notably, module eigengene (ME) corresponds to the first principal component of each module. Module membership (kME) is a measure of the ME-based intramodular connectivity, which is calculated by correlating the abundance profiles of modular members to their ME [Langfelder and Horvath, 2008, #26838]. Providing that the importance of each regulator for a functional role is determined by its degree of contribution to the module variance and by its connection strength with the other intramodular members, ranking of members according to their kME in each module (Supplemental Data Set 9) can shed light on the key or master regulator(s) for a given biological role. This approach has proven to be applicable to species with high organismic complexities such as *Homo sapiens* [Gargalovic *et al*., 2006, #11382].

Whilst each cluster was categorized and explored in detail in Supplemental Note 2, biological processes in each root zone of the two barley genotypes with module members being either induced or maintained at a high abundance level after the salt treatment are summarized in Supplemental Table 2. To validate the credibility of the global networks constructed for plants, we put one salt-induced biological process identified in Sahara, suppression of callose deposition, to the test and verified the callose abundance at four different tissue layers (focusing on epidermis, cortex, endodermis, and stele) experimentally using an immunochemical approach (Supplemental Fig. 9). In contrast to the comparable amount of callose deposited in all layers of the three root zones of Clipper after the salt treatment (Supplemental Fig. 9a,d and 9b,e and 9c,f), as deduced from the global analysis, we detected declines of callose deposition throughout the layers underneath the epidermis of Sahara in all root zones. Such declines (as indicated by the fading of orange fluorescence) were especially apparent at the plasmodesmata of cortical cell in Z3, plasmodesmata in stele and endodermis of Z2, and throughout the walls of stele and cortical cells in Z1) (Supplemental Fig. 9g,j and 9h,k and 9i,l). Further, this observation is also consistent with the fact that the *ABERRANT GROWTH AND DEATH 2* (*AGD2*), a suppressor of callose deposition [Rate and Greenberg, 2001, #18289], showed an expression pattern categorised in Module C, which is characterized by modular members with stronger salinity-induced abundance for all root-zones in Sahara than in Clipper (Fig. 7c). Altogether, these support the precision and feasibility to apply this intercorrelation approach to understand salinity responses in barley roots.

## DISCUSSION

Clipper and Sahara are two barley genotypes with known contrasting phenotypic traits in response to salt stress at an early stage of development, in which their root growth is maintained and restricted respectively [Shelden *et al*., 2013, #95836]. In this study, we investigated system-wide responses of seedling roots of the two barley genotypes to moderate salinity. We detected the spatio-temporal salt-induced perturbations to the transcriptomes, metabolomes, and lipidomes of individual root zones in each of the barley genotypes. By means of statistical over-representation of DEG and DAM (Figs. 3 and 4), we investigated the datasets from the perspective of their “extremes” and illustrated the most differential salinity responses in three different root zones of the two genotypes through integrated pathway analysis (Fig. 5; Supplemental Figs. 3 and 4). Using global co-expression correlation network analysis (Figs. 6 and 7), we approached the datasets from the perspective of “intercorrelations” among the induced pathways to demonstrate the system-wide impacts on the genotypes triggered by salinity stress (summarized in Supplemental Table 2). Through integration of the spatial and temporal omics information obtained from these approaches, we provide a novel and system-wide insight to the salt-induced modulations of apoplastic (lignin, suberin) and symplastic flows (callose) in barley roots (Fig. 8). Besides providing a comprehensive multi-omics data resource allowing deep mining of salinity-induced changes in seedlings of barley at the root zone level, we demonstrated seedling roots of different genotypes of barley could be in distinctive salinity response phases to cope with the stress, illustrating differential salt tolerance strategies could exist among the same plant species.

### Salinity-Induced Lignin Precursor Production to Sustain Clipper Root Growth

Through modification and amplification of a very limited set of core structures derived from shikimate, phenylpropanoid metabolism generates an enormous array of plant secondary metabolites ranging from monomers (such as flavonoids, isoflavonoids, coumarins, aurones, stilbenes and catechin) to polymers (such as lignins, suberins), which all share the same upstream biosynthetic steps in prior to the divergence [Vogt, 2010, #54380]. Upon short-term salt stress, our study shows that the building blocks of phenylpropanoids were diverted from the synthesis of flavonoids and suberins to the production of G- and S-units of lignins in Clipper Z2 (Fig. 5a).

Flavonoids, such as quercetin, are known as the most active and naturally-occurring inhibitors of auxin efflux carriers in a variety of plant tissues [Jacobs and Rubery, 1988, #73623]. Low levels of dihydroquercetin (Supplemental Fig. 6p), an immediate upstream precursor of quercetin, may therefore suggest the supressed inhibition of the auxin efflux carriers in Clipper Z2 compared to the same root zone in Sahara under salt stress. Operation of these carriers in turn facilitates the propagation of auxin signals along cells of Clipper Z2 and occurrence of auxin-mediated cell division and expansion. This finding is consistent with the previous phenotypic study of the two barley genotypes, in which Clipper maintains a greater root elongation rate than Sahara, even under moderate salt stress [Shelden *et al*., 2013, #95836]. This also further validates our integrated pathway analysis approach to identify their molecular differences.

Casparian strip is a specialized wall modification at endodermis, which serves as a diffusion barrier to limit apoplastic flow and re-direct solute movement back to the symplastic stream through the plasma membrane [Steudle and Peterson, 1998, #35869]. The strip is mainly constituted by rings of lignin deposited around endodermal cells and interference in lignin biosynthesis has been shown to abrogate the early strip formation in Arabidopsis [Naseer *et al*., 2012, #86004]. In monocotyledonous species, the lignin-like polymers of the Casparian strip are composed of a mixture of G- and S-units [Zeier *et al*., 1999, #45047]. In Clipper Z2, both transcript and phenylpropanoid profilings consistently show that the production and abundance of these units are significantly increased by the stress (Fig. 5a and Supplemental Fig. 6n,o). Also, an intense localization of lignins was detected at the outer stele region of the root zone after the salt treatment (Supplemental Fig. 7e,f). As in the salt tolerance mechanisms found in maize (*Zea mays* L.) [Shen *et al*., 2015, #42594], the salt-induced lignin production and its intense localization at the endodermis of Clipper Z2 likely contributed to the development of the Casparian strip closer to the root tip in response to the salt stress. There, passage of water and solutes have to undergo selective uptake via ion channels of the membranes [Apse and Blumwald, 2007, #65211]. Filtering of excessive sodium ions might therefore be achieved in Clipper Z2 by this mechanism.

For most cereal crops, deposition of suberin can be induced at cell layers such as the epidermis, outer cortex, and stele in response to salt and osmotic stresses [Schreiber *et al*., 2005, #48943]. Intriguingly, our transcriptomic data indicates that the genes involved in suberin production were down-regulated in Clipper Z2 relative to Sahara Z2 (Supplemental Fig. 5h-j). Reduced Fluorol Yellow staining in Clipper Z2 also indicated a decline in suberin levels throughout the whole root zone after salinity treatment (Supplemental Fig. 8b,h). The exodermis is a specialized outermost layer of the cortex in which Casparian strip development is inducible by salt and is found only in the wild relatives of barley, such as *Hordeum marinum* [Byrt *et al*., 2018, #25516]. In the absence of the exodermis in *Hordeum vulgare* L. genotypes such as Clipper and Sahara, low suberisation of cell layers surrounding the endodermis of Clipper Z2 would therefore imply there is no additional barrier to sodium ion entry into its root epidermis and cortex under salt stress. Consistent with this hypothesis, whole seminal roots of Clipper were shown to have higher accumulation of sodium ions than Sahara when grown under the same salinity strength [Shelden *et al*., 2013, #95836].

Plasmodesmatal conductivity is known to be regulated by the controlled build-up of callose at the plasmodesmatal neck [De Storme and Geelen, 2014, #97181]. In our study, immunochemical detection showed substantial callose deposition throughout Clipper Z2 independent of the salt stress (Supplemental Fig. 9b,e). Assuming our observed callose deposition contributed to modulating the aperture size of the symplastic channels, this may suggest a persistent restriction of symplastic flow and hence accumulation of salt in cell walls of the epidermal and cortical regions. The interwoven network of the cellulose microfibrils and pectin (such as homogalacturonan, rhamnogalacturonan I and II) is one of the major factors contributing to the cell wall strength with homogalacturonan chain interaction modulated by calcium ions [O’Neill *et al*., 2004, #69478]. Barley root cell walls have been suggested to be a “sodium ion trap” for restricting ionic movement from the root to the shoot [Flowers and Hajibagheri, 2001, #67463]. It has been shown that salt tolerant varieties have an up to two-fold greater capacity of ion adsorption than sensitive ones [Flowers and Hajibagheri, 2001, #67463], suggesting that the excessive amounts of sodium ions in the apoplast might displace calcium ions and thus weaken pectin chain calcium ion cross bridges [Ravanat and Rinaudo, 1980, #32213]. To the best of our knowledge, there is no compelling evidence to support the presence of an active exclusion mechanism for the removal of an excess of sodium ions from the apoplast of Clipper. Assuming the root cell wall was under an optimal pH required for the interaction of sodium ions and uronic acids of pectin, presence of such a high level of apoplastic sodium ions would in turn weakens the cell wall strength of roots of Clipper, implying a shortcoming of this tolerance strategy for supporting the long-term development of this genotype.

Notably, production of lignin G-units was detected in both Z2 and Z3 of Clipper, but S-unit precursors of lignin were found only in Z2, not in Z3 under the salt stress (Supplemental Fig. 6n,o). Lignin G-units are known to be a major component of tracheary elements [Higuchi, 1990, #93910], which are the key components of xylem vessels that provide mechanical resistance in plants against the negative pressure associated with the transport of minerals and water to the aerial tissues in the rising sap [Turner, 1997, #19143]. The continuance of synthesis of these units in Clipper Z3 is likely to reinforce and waterproof these cells. This suggests that the vital function for preventing the root structures from collapse and maintaining the hydro-mineral sap distribution to the whole plant served by the tracheary elements likely be independent of the salt stress.

On the whole, roots of Clipper seedlings could adopt a “growth-sustaining” strategy, which maximises root growth to increase the likelihood of overcoming the unfavourable saline conditions, but with the trade-off of developing a less effective epidermal or cortical barrier with suberin for preventing the subsequent salt accumulation in the root cortex (Fig. 8a).

### Salinity-Induced Flavonoids and Suberin Production to Shield Roots of Sahara

Unlike in Clipper Z2, our integrated pathway analysis suggests that lignin production in Z2 of Sahara was not triggered by high salinity. Instead, the building blocks of phenylpropanoids in this root zone were in part diverted to the production of flavonoids, implying suppression of root cell elongation, and in part to suberin (Fig. 5b).

Under normal growth conditions, suberization of root cells initiates mostly in the endodermis subsequent to Casparian strip formation [Geldner, 2013, #11314]. These wall modifications restrict the apoplastic uptake of water and solutes into endodermal cells. Under osmotic stress, increased numbers of suberized endodermal cells were observed at the late elongation zone of barley roots [Kreszies *et al*., 2018, #91855]. Under salinity stress, cereal crops such as maize would, however, further expand their apoplastic diffusion barrier by inducing the suberisation of cell walls in the entire root cortex in order to limit water loss from the cell layers and salt entry into xylem vessels [Andersen *et al*., 2015, #82669]. In this study, we detected high levels of suberin synthesis-related gene expression and localization of suberin throughout Z2 of Sahara, but not of Clipper (Supplemental Figs. 5h-j and 8n,t). This suggests that Sahara responds in a manner similar to other cereal crops under salt stress by hindering apoplastic transport in Z2 via suberin deposition. Consistently, our global co-expression correlation study indicates high level of salt-induced *AGD2* transcripts, a factor known for supressing callose deposition [Rate and Greenberg, 2001, #18289] (Supplemental Table 2: Sahara, AZ). Such inhibition across the subepidermal regions of Sahara Z2 (especially at the stele and endodermal regions) was confirmed by the immunochemical detection (Supplemental Fig. 9h,k). Callose deposition is known to be crucial for regulating the closure of plasmodesmata [De Storme and Geelen, 2014, #97181]. In the heavily suberised and cortical cells of Sahara Z2 with restricted apoplastic movement of nutrients taken up from the rhizosphere, inhibition of the callose deposition at plasmodesmata thus reduces the symplastic transport barrier allowing sharing and distribution of resources via the symplastic passages.

Furthermore, irrespective of the salt treatment, production of suberin (Supplemental Fig. 5h-j) and G-units of lignins (Supplemental Fig. 6n) persisted in Sahara Z3, inferring the vital importance of these precursors in the maturation of the Casparian strip and tracheary elements, respectively. Notably, unlike the untreated control, Basic Fuschin staining of Sahara Z3 showed an intense deposits of lignin around the meta- and prote-xylemic cell walls, accompanied by a small amount of lignins laid at wall of endodermis and pericycle after salt treatment (Supplemental Fig. 7c,d). In absence of the widespread salt-induced suberization of cells in epidermal and cortical layers observed in Sahara Z2 after salt stress (Supplemental Fig. 8t,s), these special arrangement of lignins at the stele of Sahara Z3 could serve as the last barriers of salt ions carried by apoplastic flows, before their entry to vasculature and be uplifted to rest of the plant parts. Further, similar to the response of the Sahara Z2, a boost in production of flavonoids was also observed in Sahara Z3 after the salt treatment (Supplemental Fig. 6p). This implies a comprehensive salt- or osmotic-induced growth restriction was triggered in both the zones of elongation (Z2) and maturation (Z3) in Sahara roots, which is consistent with the previous physiological data [Shelden *et al*., 2013, #95836]. Taken together, seedling roots of Sahara appear to implement a “salt-shielding” strategy. Such strategy restricts salt from import into the roots and minimizes water loss from root cells under the unfavourable high salinity conditions, but at the expense of the rate of growth (Fig. 8b).

### Distinctive Phases of Salinity Responses Observed in Clipper and Sahara

As defined by [Julkowska and Testerink, 2015, #28992], responses of plant cells during the exposure to salinity stress can be categorized into four main phases, namely early signalling (ES) phase, quiescent (Q) phase, recovery (R) phase, and recovery extent (RE) phase. Responses induced at the ES phase, such as the salt overly sensitive (SOS) pathway [Shi, 2002, #69407] and aquaporin internalization [Prak *et al*., 2008, #64045], can be triggered and completed within seconds or mostly hours upon exposure to salt stress [Julkowska and Testerink, 2015, #28992]. In this study, root zones of the two barley genotypes were presumably in stage of Q, R, or RE phase after three days of growth on media enriched with salt. Notably, in line with the striking growth differences observed amongst plant organs and between main and lateral roots in response to salt [Julkowska *et al*., 2014, #56802], our global co-expression correlation study reveals that salinity impacts the two barley genotypes remarkably differently in terms of the phase of responses reached by their individual root zones. Implications from the molecular and hormonal clues of the study are summarized in Supplemental Table 2 (STable2) and discussed below.

Upon exposure to salt stress, inhibition of cell cycle progression restricted the cell division and differentiation processes in Sahara Z1 (STable2: Z1, Sahara). As substantial repression of the reactive oxygen species (ROS)-scavenging mechanisms in combination with the ethylene-mediated ROS accumulation were detected in this root zone, cells in Sahara Z1 were likely retained at Q phase and not RE phase in response to the salt treatment. Notably, the ROS-related activities in the apoplast mediate cell wall stiffening through crosslinking of glycoproteins and phenolic compounds, which are known to be the milestone events detected only at the Q phase upon salt stress [Tenhaken, 2014, #31044]. By contrast, divisions of cells in Clipper Z1 were maintained and the corresponding biological processes for supporting rapid cell expansion, such as cellulose biosynthesis and cell wall loosening, were observed in Clipper Z2 (STable2: Z1, Clipper; Z2, Clipper). Although the positive modulation of cell divisions could indicate Clipper Z1 was in the stage subsequent to the Q phase (i.e. either R or RE phase), the significant upsurge of biosynthetic enzymes involved in brassinosteroid biosynthesis and initiation of the ROS-scavenging mechanism suggest Clipper Z2 was in R phase, and yet to be in RE phase. There are insufficient hormonal clues to help define the phase of responses for Clipper Z3 (STable2: Z3, Clipper). For Sahara Z2 and Z3, salt stress induced the expression of *C-REPEAT BINDING FACTOR 3* (*CBF3*) (STable2: Z2, Sahara; Z3, Sahara). In the presence of CBF, GIBBERELLIN 2-OXIDASE 7 (GA2OX7) specifically deactivates the bioactive C-20 gibberellins (GA) [Zhou *et al*., 2017, #12167]. Assuming the amount of bioactive GA was minimal under the action of GA2OX7 in barley, GA signalling and thus its growth-promoting function was restricted in response to salinity stress, implying Sahara Z2 and Z3 was retained at Q phase after the three days of salt treatment.

Strengthening the conclusion drawn from the integrated pathway analysis, our global correlation study indicates that the Z2 of Clipper proceeded to R phase for restoration of its growth rate, while all root zones of Sahara remained at a prolonged Q phase in response to the extreme salinity conditions.

Furthermore, in addition to diverting the resources for maintenance of root growth, a range of known downstream salt tolerance mechanisms, such as polyamine transport and toxin catabolism [Frommer *et al*., 1995, #20776; Roxas *et al*., 1997, #10648], were also activated in Z2 of Clipper in order to cope with the salinity stress (STable2: Z2, Clipper). Notably, the majority of the tolerance mechanisms triggered were different between Z2 and Z3 of Clipper, where biological processes including seed oil body formation, glucosinolate hydrolysis, and nicotianamine biosynthesis [Bonneau *et al*., 2016, #34836; Eriksson *et al*., 2002, #67886; Shimada *et al*., 2008, #93537] were either induced or maintained in Z3 of Clipper, but not in Z2 (STable2: Z3, Clipper). Only hydrolysis of glucosides [Markham *et al*., 1998, #28207] has widespread up-regulation in all root zones of Clipper (STable2: AZ, Clipper), suggesting the salt tolerance strategies adopted by this genotype are mostly root zone-dependent; a mechanism that can only be explicitly revealed by the spatial multi-omics approach described here.

In contrast, with only two salt-induced biological processes, membrane steroid modulation and inhibition of cell cycle progression, with members up-regulated or maintained at high abundance in Sahara Z1 (STable2: Z1, Sahara), seven out of seventeen processes in Sahara were shared among two root zones (STable2: asterisks). Members involved in the eight processes remained such as biosynthesis of glycine betaine, modulation of GA signalling, and LTP-mediated tolerance response in all root zones of Sahara were found to be induced or maintained at higher abundance than in Clipper (STable2: AZ, Sahara). This finding suggests the tolerance mechanisms triggered in Sahara were mostly root zone-independent. Such independence is also consistent with our conclusion that all root zones of Sahara are in the same phase of the salinity response.

### Perspectives

While single-omic analyses identifies candidates and processes that are fragmentary and disconnected, in this study, we show that the integrative spatial multi-omics approach can integrate the molecular changes detected at levels of transcriptomes, metabolomes, and lipidomes to provide novel systematic insights into the early salt tolerance strategies in barley. By considering the datasets from the perspective of “extremes”, we demonstrate that Clipper could adopt a “growth-sustaining” strategy to increase the likelihood of escaping from adversity by root growth but may require a trade-off of developing a less effective barrier against subsequent salt accumulation in roots. In contrast, the data suggest that Sahara adopts a “salt-shielding” strategy to block out salt access to the interior of seedling roots, but at the expense of growth rate upon salt stress. Furthermore, by considering the datasets from the perspective of “intercorrelations”, we proposed that two distinctive salt tolerance mechanisms could exist in different genotypes of barley, in which the mechanisms were growth-oriented and root zone-dependent in Clipper but were more salt tolerance-oriented and root zone-independent in Sahara. Understanding the differing natural strategies adopted by the barley genotypes may help in designing plants to cope with the predicted increase in salinity stress, which will impact our ability to maintain yield in important food and feed crops in future.

## METHODS

### Plant Materials

Genotypes of barley (*Hordeum vulgare* L.) were originally sourced from the Australian Centre for Plant Functional Genomics at the University of Adelaide. Two genotypes of barley, the domesticated malting cultivar Clipper (Australia) and the landrace Sahara 3771 (North Africa), were used for the omics analyses and phenylpropanoid detection in this study, and were selected based on previously reported physiological diversity in salt tolerance [Shelden *et al*., 2013, #95836; Widodo *et al*., 2009, #52958].

### Growth Conditions and Sample Preparation

The experiment and sample collection were described previously [Hill *et al*., 2016, #53060]. In short, uniformly sized barley seeds were grown under control (nutrient medium without additional NaCl) and salt-treated (nutrient medium supplemented with 100mM NaCl) conditions. Seminal roots were dissected after three days of growth on agar media, whereby a 1.5 mm long section marked ‘Z1’ (meristematic zone) was taken from the root tip, a second section marked ‘Z2’ (elongation zone) was dissected up to the third section, ‘Z3’ (maturation zone), which was excised at the point of visible root hair elongation. For study of the transcriptomes (RNA-seq) and primary metabolomes (GC-QqQ-MS sugar and organic acid quantification, LC-MS amine quantification, GC-Q-MS FAME quantification, and lipid analysis), four biological replicates were generated for each sample in four separate experiments totalling 48 samples. For detection of the phenylpropanoids, three biological replicates were prepared for each sample in three independent experiments with a total of 36 samples. All dissected seminal roots were collected into pre-chilled 1.5 mL tubes, immediately snap-frozen in liquid nitrogen, weighed, and then stored at −80 °C until extraction of RNA, primary metabolites, lipids, and phenylpropanoids.

### Functional Annotation of the New Barley Reference Genome

To further enrich the functional annotations of the mapping base for RNA-seq, the latest version of the new barley reference genome sequences (cv. Morex v2) and the genome structural annotation files were obtained from IPK Barley server of the International Barley Sequencing Consortium (IBSC) [Mascher *et al*., 2017, #46618]. The total population of coding sequences of the genome was extracted by the gffread utility of Cufflinks [Trapnell *et al*., 2012, #90292] and refined using the degapseq script of EMBOSS 6.6.0.0 [Rice *et al*., 2000, #83873]. The latest version of Basic Local Alignment Search Tool (BLAST) was obtained from the FTP server of the National Center for Biotechnology Information (NCBI) [Altschul *et al*., 1990, #64088], and a local BLAST pipeline was constructed in eight NeCTAR Research Cloud instances in Ubuntu 16.04 LTS (Xenial) environment [Li *et al*., 2018, #18413]. The total population of translated coding sequences of the barley genome were BLASTx searched against three protein sequence databases, i.e. TAIR10 [Lamesch *et al*., 2012, #34604], UniProtKB/Swiss-Prot [The UniProt Consortium, 2017, #70057], RAP-DB [Sakai *et al*., 2013, #15802], and two ontology databases i.e. Gene Ontology (GO) [Ashburner et al., 2000, #93070] and KEGG Ontology (KO) [Kanehisa and Goto, 2000, #77796]. The latest version of InterProScan-5 and Panther models 10.0 were obtained from the FTP server of the European Bioinformatics Institute (EMBL-EBI) and the getorf script of EMBOSS was applied to make InterProScan-5 to be compatible to nucleotide inputs. Scanning of InterPro protein domains databases was performed according to the user manual [Jones *et al*., 2014, #35110]. Only the top hits of each coding sequence with the lowest e-values were listed in the functional annotation list and considered for biological interpretation.

### RNA Isolation, Sequencing, Read Processing, and Mapping

RNA isolation and sequencing were described previously [Hill *et al*., 2016, #53060]. In short, the total RNA was extracted from 50 mg root tissue separately per genotype, treatment, and root zone using the Qiagen RNeasy kit following the manufacturer’s protocol. All RNA-seq libraries were constructed and paired-end sequenced (100bp) on an Illumina Hi-Seq 2000 system at the Australian Genome Research Facility (Melbourne, Australia). Four lanes were used for each genotype, and all 48 samples were run on a single flow cell. The RNA was sequenced to a depth of approximately 31 million read-pairs per sample per lane, giving a total of 1.48 billion reads (749 million read-pairs).

Paired-end libraries of raw reads from the RNA-seq were verified and converted using FASTQ Groomer [Blankenberg *et al*., 2010, #19632] and sequence quality was validated using FastQC [Andrews, #9190]. Based on the outcomes of the read quality assessment, threshold was defined (q=20; minimum read length: 24; Illumina TruSeq Adaptor primers removed; singletons discarded) and Trimmomatic was applied to trim reads for quality [Bolger *et al*., 2014, #30435]. Mapping or paired-read alignment was performed via HISAT2 [Kim *et al*., 2015, #48577] and the sorted BAM files were subjected to HTSeq code [Anders *et al*., 2015, #86271] for generation of the counting matrix using the genome structural annotation available from IBSC [Mascher *et al*., 2017, #46618].

### DEG Determination and Enrichment Analysis of Gene Ontologies

To prepare for DEG determination, we filtered the lowly expressed genes from the matrix were filtered based on a minimum CPM threshold of 11.5 present in at least four samples, which corresponds to an average read count of 10-15 across the 192 libraries, to minimise the multiple testing burden when estimating false discovery rates [Robinson *et al*., 2010, #64898]. TMM normalization was applied to the transformed CPM matrix to eliminate composition biases between libraries [Robinson and Oshlack, 2010, #78960]. Multidimensional scaling of the TMM-normalized matrix explicitly revealed one biological replicate of Clipper control at Z3 as an outlier and was therefore excluded from all subsequent analyses. Variation of library sizes, sample-specific quality weighting, and mean-variance dependence of the data matrix were addressed by the voom transformation workflow available in limma package (v.3.7) [Ritchie *et al*., 2015, #41043]. Detailed procedures for estimating group mean and gene-wise variances, as well as fitting of basic and interaction GLM to test for differential expression were detailed in [Smyth *et al*., 2002, #61553]. Notably, as discussed by [Zhang and Cao, 2009, #91391], assumptions required for fold-change filtering and *t*-statistic adopted in DEG determination were contradictory, therefore only the *t*-statistic-based adjusted *p* value was applied as a cutoff in this study.

For enrichment analysis of GO, BiNGO was applied to determine the overrepresented GO terms in each DEG list focusing only on the GO Biological Processes category [Maere *et al*., 2005, #47202]. Unless otherwise specified, the analyses were performed using the hypergeometric test with the whole barley annotation as a reference set and Benjamini-Hochberg FDR correction with *q* value cutoff at 0.05. Each enrichment list was summarized by REVIGO with small (0.5) allowed similarity [Supek *et al*., 2011, #81163] and enrichment networks resulted were visualized in Cytoscape (v.3.4.0) [Shannon, 2003, #25599].

### Metabolite and Lipid Quantification

Metabolites (sugars, organic acids) were quantified as described in [Dias *et al*., 2015, #62168]. Amines and amino acids were quantified as described in [Boughton *et al*., 2011, #52434]. Fatty acids were quantified as described in [Eder, 1995, #31243]. Lipids were quantified as described in [Natera *et al*., 2016, #68786]. Phenylpropanoids were extracted from three biological replicates of root tissues (10mg) per genotype, treatment, and root zone from exactly the same growth settings, using 500 µl of cold methanol each. After homogenisation by CryoMill (RETSCH), samples were agitated for 15 min at 70°C with 10,000 rpm in a thermoshaker (Eppendorf), then allowed to cool down the extract before centrifuged for 5 minutes at room temperature with 14,000 rpm. Supernatant was transferred into a clean Eppendorf tube for further clean-up process using solid phase extraction (SPE) cartridges. For SPE clean-up process, 60 mg Agilent Bond Elute Plexa cartridges were conditioned using 1 mL of methanol, followed by 1 mL of water. The supernatant from root extract was loaded and washed by passing 1 mL of methanol, then metabolites were eluted using 400 µL of methanol, followed by 400µL of 5% formic acid in methanol. Combined elute was dried down in a speed vacuum and reconstituted in 100µL of 50% methanol: water prior to LC/MS analysis.

Phenylpropanoids were separated by an Agilent 6490 triple quadrupole mass spectrometer coupled to an Ultra High-Performance Liquid Chromatography (LC-QqQ-MS) (Santa Clara, CA, USA). An Agilent luna C18 column (2.1 mm x 150 mm, 3µm) was used for compound separation. The mobile phase composition included A: 10mM ammonium acetate in methanol/ water/ Acetonitrile (10/85/5, v/v/v) and B: 10mM ammonium acetate in methanol/ water/ acetonitrile (85/10/5, v/v/v) with a gradient elution: 0-10 min. 45% A; 10-20 min, 55% to 100% B; 20-22 min; 100% B; 22-25 min; 55% B to equilibrate the column to initial conditions. The flow rate of the mobile phase was maintained at 0.2 ml min^-1^ and the column temperature was maintained at 50°C. The needle wash was 20% (v/v) acetonitrile in water with sample injection volume of 5 µl. Analysis was performed using Agilent MassHunter acquisition software, version 7. Compounds were quantified based on calibration curves prepared using authentic standards.

Mass spectrometry detection was performed using ESI source operated in positive ion mode. The source parameters were set as; capillary voltage 4.0 kV, iFunnel high pressure RF in positive and negative mode at 130V, low pressure RF in positive and negative mode at 60V; source temperature 200 °C, sheath gas temperature 400 C°, gas flow 12 L min^-1^, sheath gas flow 12 L min^-1^, fragmentor voltage 380 V and cell accelerator 5V. Data was collected using in-house multiple reaction monitoring (MRM) developed based on individual standards. Dwell time for each compound was set as 10 ms and data was quantified using MassHunter Quant software version 7.

### DAM Determination and Metabolite Set Enrichment Analysis

Data matrices correspond to each type of primary metabolomes and phenylpropanoids were standardized by sample weights to achieve unit-conformity across different extraction and detection workflows. To reduce systemic bias during sample collection and impact of the large feature (metabolite) values, log-transformed matrices were normalized by median across samples and mean-centred, respectively [van den Berg *et al*., 2006, #85175]. Each normalized matrix was individually evaluated for unwanted variances by means of relative log adjustment - within group (RLAwg), principal component analysis (PCA), and hierarchical-clustering (HCR) [Xia and Wishart, 2011, #43779], which unambiguously indicated one out of four of the biological replicates in the primary metabolome detection as an outlier which was therefore excluded from all subsequent analyses. Potential batch effects attributed to sample degradation and/or instrumentation platform differences were evaluated and adjusted using the RUV-R method [Livera *et al*., 2015, #4908]. For determination of DAM, a limma-based linear modelling algorithm fitted with moderate statistics (simple Bayesian model) developed by [Livera and Bowne, 2014, #14825] was adopted to construct the basic and interaction GLM contrasts required for determination of DAM. MBROLE (v.2.0) with use of the full database as reference set, but selected only the functional roles that are non-ambiguous and can be found in the *Plantae*, were utilized to detect the enrichment of metabolite sets of each list of DAM [López-Ibáñez *et al*., 2016, #73627].

### Integrated Pathway Analysis

To integrate the omics datasets at the pathway level, coding sequences of DEGs identified from the differential analyses of the twelve transcriptomes upon salt treatment were translated and BLASTx searched against the *Arabidopsis* genome release (TAIR10, version: Jun 2016) and KEGG pathway repository (version: May 2017). Only matches with E-value < 1.00E-4 (or smallest possible E-value in the case of multiple hits for the same gene) against either or both of the databases were retained and corresponding K numbers in the KEGG repository were fetched for the subsequent integration step. For primary metabolites, the C numbers of DAMs detected in each LC/GC-MS-based quantification were determined by comparison of their chemical structures, formulae, molecular weights, and/or IUPAC nomenclatures between the reference standards used and the KEGG compound repository (May 2017). KEGG mapping of the K and C numbers acquired was performed against the pathway repository of *Arabidopsis thaliana*, which is the most comprehensive and representative pathway collection among all plant species within the KEGG database, following the procedures as stated previously [Aoki and Kanehisa, 2005, #40791]. Generic outputs from the KEGG mapper (including: ath01100 Metabolic pathways, ath01110 Biosynthesis of secondary metabolites) were defined as outputs from the KEGG mapper common to any kind of inputs and were therefore excluded from the ranking process. Only pathways statistically enriched in terms of GO categories (as determined by BiNGO) and of metabolite sets (as determined by MBROLE2) were ranked in descending order according to the number of significant DEG and DAM matches.

### Correlation Network

Abundance matrices of the total population of DEG and DAM from each barley genotype were concatenated as individual inputs for the Weighted Correlation Network Analysis (WGCNA). Processing of the matrices and network comparisons were performed as described [Langfelder and Horvath, 2008, #26838]. In brief, matrices were evaluated for missing value using the goodSamplesGenes function and any outliers were determined by hierarchical clustering. Scale-free topology and mean connectivity of each network were plotted against the soft thresholding power to derive the optimal adjacency or dissimilarity. Two coexpression correlation networks (also known as hierarchical clustering of transcript and metabolite abundance) specific to Clipper and Sahara were built based on dissimilarity-based topological overlap matrix (TOM). Modules of each network were defined by dynamicTreeCut and modules unique to each network were determined the matchModules function. Comparability of the two matrices was confirmed by verifying the correlation of ranked expression and ranked connectivity between the two datasets. Module preservation between the independent coexpression-correlation networks of Clipper (as ‘reference’ set) and Sahara (as ‘test’ set) were calculated by the ‘modulePreservation’ function of the WGCNA package v1.61, which outputted the ‘Zsummary.pres’ value for each module based on preservation-statistics and module quality-statistics (including quality, preservation, accuracy, reference separability, and test separability). Z>10 (including modules brown, turquoise, yellow, blue, greenyellow, and green), 5<Z<10 (including black, purple, red, cyan, pink), and Z<5 (including magenta, tan, salmon) indicate high preservation, moderate preservation, and low preservation or modules with significant contrast, respectively.

Modules with Z-score <10, excluding module ‘tan’, which was determined as noise, are defined as weakly preserved modules or modules with significant contrast between the two barley genotypes. Parallel plots for showing either positive or negative correlation of different abundance clusters (within 99^th^ percentile) were generated using the ggplot package of R software. The most representative trend or centroid of each module represented by purple solid lines was determined by k-mean clustering (distance method: Pearson) with optimal number of clusters calculated using the within-group sum of square method [Madsen and Browning, 2009, #85081]. Module memberships (kME) of genes and metabolites harboured among module or cluster unique to either network or significantly different to the other network were calculated by signedKME function of the WGCNA package.

### Histochemical and Immunochemical Microscopy

Roots of barley cultivars Clipper and Sahara, grown on agar media supplemented with either 0 mM or 100 mM NaCl for three days, were fixed in 4% paraformaldehyde overnight at 4°C and then washed in phosphate-buffered saline (PBS). For lignin and suberin staining, the roots were embedded in 6% agar followed by sectioning of 80 µm thick sections using a VT1000 S vibratome (Leica Microsystems). Sections for lignin staining were cleared using Clearsee [Kurihara *et al*., 2015, #40262] and stained using 0.2% w/v Basic Fuchsin and 0.1% w/v Calcofluor White (general cell wall stain) [Ursache *et al*., 2018, #46620]. Vibratome sections for suberin staining were placed in 0.01% w/v Fluorol Yellow 088 (Santa Cruz Biotechnology) in polyethylene glycol 200 for 1 h at 90°C [Brundrett *et al*., 1991, #90976] followed by counterstaining with 0.5% aniline blue for 30 mins. Roots for callose labelling were dehydrated in an ethanol series followed by infiltration and embedding in London White Resin (LRW) (ProSciTech). Sections (1 µm) were cut using an Ultracut S ultramicrotome (Leica Microsystems) and labelled with the primary (1:3)-β-glucan antibody (Biosupplies Australia) [Meikle *et al*., 1991, #4671] at a concentration of 1:300 [Wilson *et al*., 2015, #56414] followed by the secondary anti-mouse 568 Alexa Fluor antibody (Thermofisher) at a 1:200 dilution.

A Nikon C2 confocal microscope (Coherent Scientific, Australia) equipped with a spectral detector was used to image the cell wall fluorescence using the following settings: Basic Fuchsin - ex 561 nm, em 600-650 nm; Calcofluor White – ex 405 nm, em 425 - 475 nm; anti-mouse Alexa Fluor 568 antibody – ex 561 nm, em 570-650 nm. Fluorol Yellow staining was imaged using a Leica DM6000 microscope equipped with a Leica DFC 450 camera using the I3 (GFP/FITC) filter. Images were analysed using FIJI (NIH).

### Accession Number

Raw reads of RNA-seq applied in this work were deposited in the ArrayExpress database (http://www.ebi.ac.uk/arrayexpress) under accession number E-MTAB-4634. Sequence information for all Arabidopsis genes described in this article can be found in TAIR (www.arabidopsis.org) using the following accession numbers: *CEL1* (AT1G70710), *GLR2.7* (AT2G29120), *EXPB2* (AT1G65680), *AAT1* (AT4G21120), *GSTU18* (AT1G10360), *CESA1* (AT4G32410), *CESA3* (AT5G05170), *EXPA11* (AT1G20190), *EXPB4* (AT2G45110), *EXPA7* (AT1G12560), *DET2* (AT2G38050), *TCP8* (AT1G58100), *TCP15* (AT1G69690), *TCP23* (AT1G35560), *XTH13* (AT5G57540), *DSEL* (AT4G18550), *OLE1* (AT4G25140), *EXPA2* (AT5G05290), *EXPA13* (AT3G03220), *EXPB2* (AT1G65680), *FMO1* (AT1G19250), *MBP1* (AT1G52040), *CBF3* (AT4G25480), *GA2OX7* (AT1G50960), *DALL1* (AT4G16820), *NAS3* (AT1G09240), *NAS4* (AT1G56430), *ACX3* (AT1G06290), *SIP4* (AT2G30360), *APX1* (AT1G07890), *PEAMT* (AT3G18000), *ADH10A8* (AT1G74920), *GID1C* (AT5G27320), *AGD2* (AT1G60860), *XTH20* (AT5G48070), *ESK1* (AT3G55990), *PAE7* (AT4G19410), *MUCI10* (AT2G22900), *PGSIP1* (AT3G18660), *DIR1* (AT5G48485), *AZI1* (AT4G12470), *PR-4* (AT3G04720), *RAP2.11* (AT5G19790), *CALS3* (AT5G13000), *CALS7* (AT1G06490), *SOX* (AT3G01910), *CYP51A2* (AT1G11680), *CDC2* (AT3G48750), CYCP4;1 (AT2G44740), *IBR5* (AT2G04550), *GAE* (AT4G30440), *EARLI1* (AT4G12480), and *STE1* (AT3G02580).

## ACKNOWLEDGMENT

The authors would like to thank Ms Nirupama Jayasinghe (Metabolomics Australia, School of BioSciences, University of Melbourne) for providing excellent technical assistance for GC-QqQ-MS and LC-QqQ-MS analyses, Mr Daniel Sarabia and Mrs Cheong Bo Eng (School of BioSciences, University of Melbourne) for root sample collection for phenylpropanoid detection, as well as technical staff at the Australian Genome Research Facility (AGRF, Melbourne) for next-generation sequencing. The authors are grateful to Melbourne Bioinformatics (www.melbournebioinformatics.org.au) for access to the HPC and data storage facilities, and to the Victorian Node of Metabolomics Australia, which is funded through Bioplatforms Australia Pty. Ltd., a National Collaborative Research Infrastructure Strategy (NCRIS), 5.1 Biomolecular platforms and informatics investment, and co-investment from the Victorian State Government and The University of Melbourne. We would also like to acknowledge the provision of facilities from the Biosciences Microscopy Unit (BMU-05) and Ms Ewa Anna Reda for her pilot experiments on root sectioning and lignin staining. This research was supported by use of the Nectar Research Cloud, a collaborative Australian research platform supported by the Australian Government through the National Collaborative Research Infrastructure Strategy (NCRIS). UR is grateful to the Australian Research Council for funding this work through a Future Fellowship award.

## AUTHOR CONTRIBUTIONS

C.B.H., M.S.D., M.C.S, A.B., and U.R. designed the experimental part of the research. C.B.H. performed the salinity experiment, collected and extracted samples, processed MS data, and performed initial metabolite data analyses. T.R. carried out the phenylpropanoid detection and processing of mass spectrometry data. A.V.D.M. performed histological work on barley root sections. W.W.H.H. designed and implemented the bioinformatics part of the research, and performed all subsequent statistical and computational analyses, including functional annotation of barley genome with HPC, GLM-based differential analyses, and omics data integration via integrated pathway analysis and global expression correlation networks. W.W.H.H., C.B.H, M.S.D., M.C.S, A.B., and U.R. interpreted the data and wrote the article. All authors revised, edited and approved the manuscript.

